# Mechanical Properties of DNA Replication

**DOI:** 10.1101/699009

**Authors:** Stuart A. Sevier

**Affiliations:** Center for Theoretical Biological Physics, Rice University, Houston, TX 77005, U.S.A. and Institute for Medical Engineering and Science, Massachusetts Institute of Technology, Cambridge, MA 02139, USA

## Abstract

Central to the function of cellular life is the reading, storage and replication of DNA. Due to the helical structure of DNA, a complicated topological braiding of new strands follows the duplication of the old strands. Even though this was discovered over 60 years ago, the mathematical and physical questions this presents have largely gone unaddressed. In this letter we construct a simple idealized model of DNA replication using only the most basic mathematical and mechanical elements of DNA replication. The aim of this is to reveal the mechanical balance of braided, replicated DNA against the twist of unreplicated DNA at the heart of the replication process. The addition of topoisomerase action is included presenting a balancing force offering a glimpse into the ways in which cells maintain this balance. Additionally the physical basis for recently observed replication/replication and replication/transcription conflicts are examined showing how gene orientation and size can impact DNA replication.

---

For life to continue cells must create new versions of themselves. A central task in this process is the creation and proper segregation of a new copy of DNA. This must be accomplished while simultaneously leaving DNA available for important cellular functions. Soon after the helical nature of DNA was uncovered [1] it was realized that each parent strand serves as a template for two daughter strands [2] (deemed semi-conservative replication). This results in two daughter strands that are, without topological breaks, braided around each other. This braided state presents a major topological and physical barrier to cellular replication and function which must be undone before division can take place [3, 4]. The ability for cells to undo this braided state, while undergoing normal functions, presents central competition in DNA replication which must be balanced against topoismerase action and transcription for cells to successfully divide.

The competition may underlay many important aspects of DNA replication such as non-local effects and coordination in DNA damage, replication fork conflicts and transcriptional interference [5–9]. Consequentially, understanding the mathematical and physical nature of DNA replication is of upmost important in understanding many aspects cellular function. Many biological aspects of DNA replication and function of been uncovered [5, 10, 11]. However, a physical model of this simple process has not been developed leaving many basic questions concerning the mechanical competition present during replication unanswered [10]. The answer to these questions may offer solutions to a number of emerging issues concerning cellular function.

In this letter we construct a simple picture of the physical, mechanical process of DNA replication. In order to try and reveal the central elements of the topological competition present during DNA replication we we will ignore the differences between different organisms. We attempt to take some of the first steps in addressing this competition and it’s resolution by studying a simple idealized mode of DNA replication with the topological constraints of the most basic version of replication as central features. To do this we will divide the system into replicated and unreplicated regions and focus on the resulting DNA braiding, which occurs in the replicated region, and the DNA super-coiling which occurs in the unreplicated region (see figure 1).

**FIG. 1:**
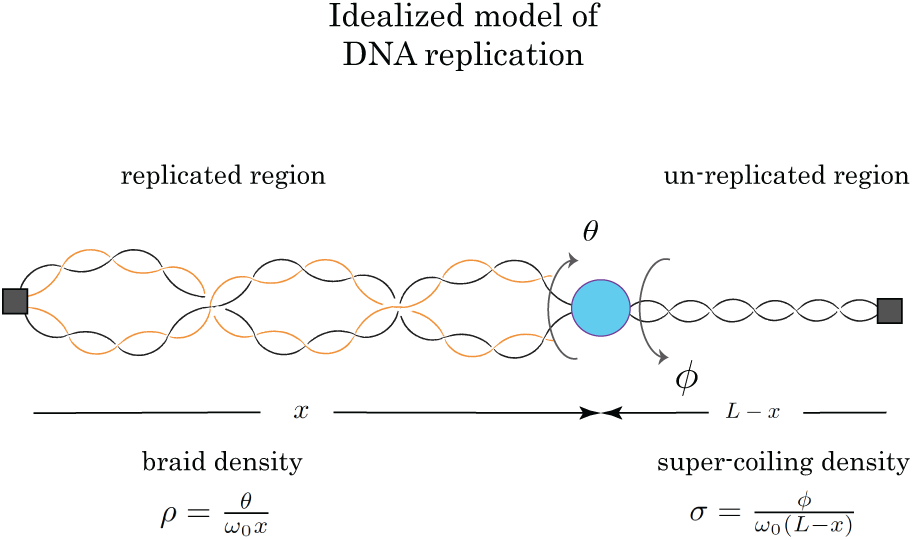
A cartoon illustrating an idealized model of DNA replication. Fork progression shown at a distance *x* from the start of replication results in braided replicated DNA strands rotated around each other *θ* times and super-coiled un-replicated DNA over-rotated *ϕ* times. The two regions are separated by idealized free rotating replication machinery shown in blue. The two ends are shown as simple topological barriers (gray squares) which prohibit the free rotation of the DNA thus conserving the linking number of the parental DNA. This results in a mechanical and topological connection between *θ* and *ϕ*.

In this idealized model, the basic coordinates are the replication fork distance from the replication start site *x*, the number of times the replicated strands are wrapped around each other *θ* and the over-twisting *ϕ* of the unreplicated region (fig.1). We will take *θ* and *ϕ* to be of opposite sign so that they are both positive quantities as a function of fork position. Due to the helical nature of DNA and the semi-conservative nature of replication, these quantities must satisfy the basic topological constraint during replication when no strand passages have occurred

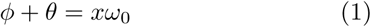

where *ω*_0_ = 1.85*nm*^*-*1^ encodes the natural linking number of DNA. As the replication fork progresses, the relative difficulty in twisting the unreplicated DNA or braiding the replicated strands determines the form of the functions *ϕ*(*x*), *θ*(*x*).The extra angle of DNA twisting *ϕ* and braiding *θ* can be determined by the balance between DNA torque of the replicated braided DNA *τ*_*rep*_ and unreplicated twisted DNA *τ*_*unrep*_ with specified boundary conditions. We will not explicitly incorporate the mechanics of the replication machinery but will comment on possible implications of its mechanical regulation.

To simplify the system we will introduce an explicit topological barrier at the point of replication origination and termination (a distance *L* from the start site) as shown in figure 1. These idealized barriers prevent free rotation of the DNA as well as linking number conservation of the original parental DNA strands. A static barrier of this nature can be explicitly introduced in an in-vitro setting and may have natural or alternative forms in vivo (discussed later). This boundary condition leads to a description of replicated DNA behavior through the replicated braid density 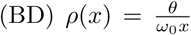 and unreplicated super-coiling density 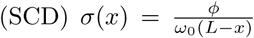. In principle these two quantities are not constant in the two regions. However, here we will only consider the case of the average, constant braid and super-coiling density neglecting transport effects. Directly substituting the BD and SCDs into equation 1 yields a topological constraint on the densities

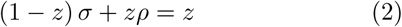

where the expression depends only on the fraction of the strand replicated 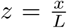. For the case of constant braid and super-coiling density, equation 2 assumes that twisting strain at the point of replication immediately spreads throughout the specified DNA length resulting in uniform braid and super-coiling density. We will neglect the dynamical response of both the braided as well as the twisted region and instead adopt the equilibrium torsional response of the two regions, resulting in constant braid and SC densities (as outlined above). We will adopt this perspective in our simple model due to DNA mechanical responses occurring on a sub-second time scale [12, 13]. A more general framework would take into account the dynamics and resulting inhomogeneity of both BD and SCD. Future work to address both of the issues will allow for insight into in-vivo experiments. However, the basic mechanical and topological constraints outlined above must still hold. These assumptions lead to a torsional constraint between the replicated and replicated regions

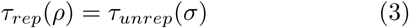

This assumption, combined with the topological constraints in equation 2 form the foundation for the most basic mechanical properties of DNA replication which will be examined in this letter. The above torsional constraint must hold no matter the details of the replication machinery, provided it is freely rotating (which is expected given current knowledge of replication machinery operation) and the only torsional constraints exist at the two ends. The determination of the BD *ρ* and SCD *σ* is made through evaluating *τ*_*rep*_(*ρ*) and *τ*_*unrep*_(*σ*) during fork progression.

In the unreplicated region the torsional response is due to the over-twisting of the single parental double stranded DNA. For super-coiled DNA at constant force its torsional response is given by an unbuckled elastic response followed by a buckled with a constant torque [14]

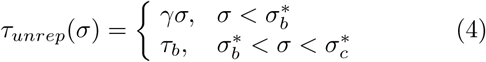

where *γ, τ*_*b*_ are the elastic torsional coefficients and buckling torque, respectively and 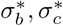 the SCDs at which buckling and a plectonemic collapse occur. The elastic response is on the order of *γ* ≈ 10^2^*pNnm* and the force dependence described in [14].

In the replicated region the torsional response is due to the braiding of the two replicated double DNA strands. Since the common picture of DNA semi-conservative replication involves freely rotating individual DNA strands, we will not consider the role of super-coiling. Additionally, because the torsion required to buckle braided DNA is higher than the constant torsion given after the unreplicated buckling transition *τ*_*b*_, we will not consider buckling in the braided, replicated strands. For the braided DNA to undergo buckling the unreplicated DNA must be in a completely plectonemic state, resulting in a collapsed replication fork, a state which falls outside our framework of understanding DNA replication. The point of buckling 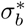, and of collapse 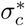,will be central to our analysis and their identification is one of the primary results of this letter. Thus, we will only consider the torsional response associated with of the braiding the two strands of DNA (due to the bending energy)[15] resulting in non-linear response to braid density

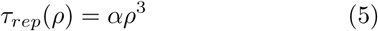

with no force dependant quantities and *α* ≈ 10^4^*pNnm* (see SM for derivation). The mechanical coefficients for both SC and braided DNA depend on temperature and physiological conditions [14, 15] (see S.M. for details) which will be fixed at the given values within this letter.

Thus, using the torsional responses given by eqs.4, 5 combined with eq.3 yields the constraint

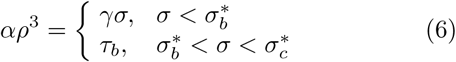

which, through the use of the topological constrain on densities (eq.2), can be used to obtain an equation explicitly for the braid density before buckling

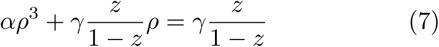

while after buckling *ρ* is given by the second line of equation 6. Since *α* ≫ *γ* and *ρ* ≪ 1 we will drop the linear term (full solution given in SM) to find the braid density as a function of replication position in the unbuckled and buckled regimes

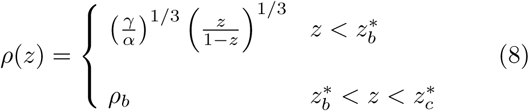

where the constant braid density *ρ*_*b*_ = (*τ*_*b*_*/α*)^1*/*3^ is given by the buckling torque in the unreplicated DNA. Using eq. 2 we can obtain the SCD in the unreplicated region

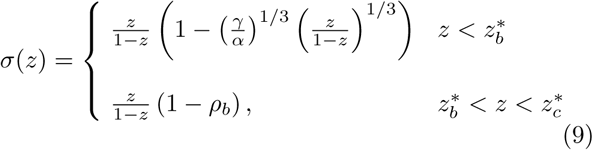

then obtaining the point at which both buckling and collapse occur respectively as

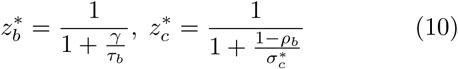

where the forms of *τ*_*b*_ and 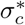 are given in [14]. The torsional response as a function of DNA replication is given by equation 2 yielding

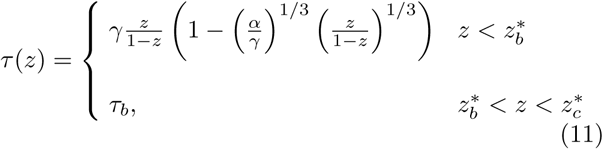

So that for *x* ≪ *L* the braid density increases as 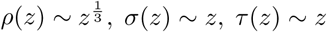 (see fig.2). As illustrated in figure 2, only a small fraction of DNA replication is possible before the replication fork enters a collapsed plectonemic state at 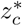.

**FIG. 2:**
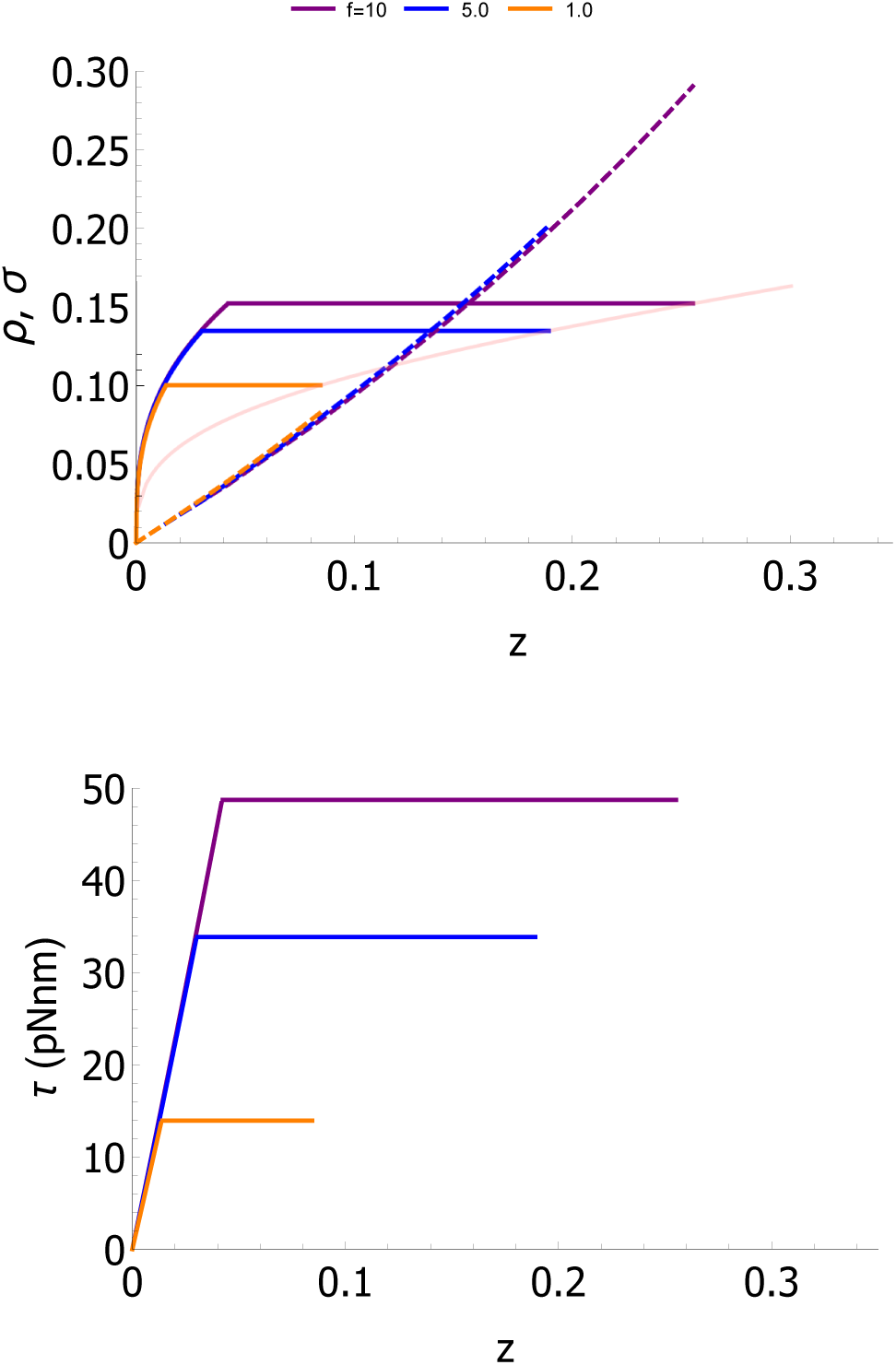
Braid density *ρ*, super-coiling density *σ* (a) and torque *τ* (b) as a function of fork position *z* = *x/L* at variance fixed forces (shown in top legend). Pink dashed curve shows collapse position *zc* for increased force as shown in figure 3. As the force in increased both the buckling *z*_*b*_ and collapse *z*_*c*_ increase but at the cost of increased buckling torque *τ*_*b*_.

Increasing 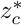 is possible by increasing the force in the system (shown by green dashed line) however a price is paid in the buckled torque *τ*_*b*_ (see fig.3) introducing a competition between the constraints placed on the amount of replication fraction possible *z* and the torque experienced during replication *τ*_*b*_. This is important, as mechanical limits to the ability of the replication machinery to unwind the unreplicated DNA may be occur before the fork reaches 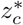 thus placing additional constraints on the mechanical limits of DNA replication. Additionally, if the replication fork was able to reach the end of the system there exists a divergence *z* →1 in the SCD. This is due to our formulation of the system in terms of densities which is ill-defined for the SCD at *z* = 1 making our approach invalid at the final point of replication completion.

To understand how DNA replication may avoid collapse or high torsion we must incorporate the action of topoisomerase is a class of molecular complexes capable of altering the linking number of DNA through strand breaks and passages [10]. In our simplified model this will be done through a dynamical equation for SCD which accounts for the generation of SCD during replication and its removal with topoisomerase. Using equation 2 we can generate the dynamical contribution of SCD due to replication fork progression

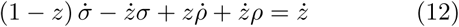

which can be used to described the dynamical state of the SCD in both the buckled and unbuckled state through the use of equation 6. For the sake of simplicity and since we are interested in how topoisomerase action can prevent an replication collapse we will consider the case of constant *ρ*, given in the buckled phase, with a constant fork velocity *z*(*t*) = *vt/L* to find

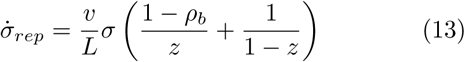

and will incorporate the action of topoisomerase with the addition of a a simple rate of removal 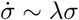. We will not specify the direct mechanisms of topoisomerase action in an effort to identify the basic elements of this dynamic competition. This yields a dynamical equation for the SCD

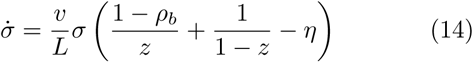

where 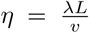 and we have used equation 2 to write the expression its seperable form. An additional term 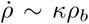, as well as alternative terms for topoisomerase removal and SCD dependant velocity, can be included in this framework but will not be examined here. This equation can be directly integrated to find the solution

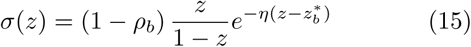

where we have used the boundary condition 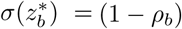 to account for matching at the buckling transition. For a collapse to be avoided we must have 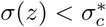 which puts a condition on the relative rates of replication *v* and topoisomerase action *λ* as well as the buckled density *ρ*_*b*_. A local maxim can occur for *σ*(*z*) at

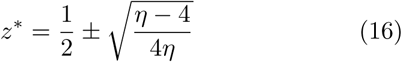

if the rates satisfy the inequality *λ* > 4*v/L*. Thus it is possible, if *ρ*_*b*_ is such that 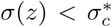 for replication to occur over the entire system, as shown in figures 3 and 4. While not a complete description of the competition between DNA SCD accumulation and removal during replication, this basic result helps uncover some of the essential properties of DNA replication which must occur for DNA to be successfully duplicated. These results point to the importance of quick topoisomerase action during replication and offer insight into the importance of extra proteins used to stabilize plectonemes during replication [16].

**FIG. 3:**
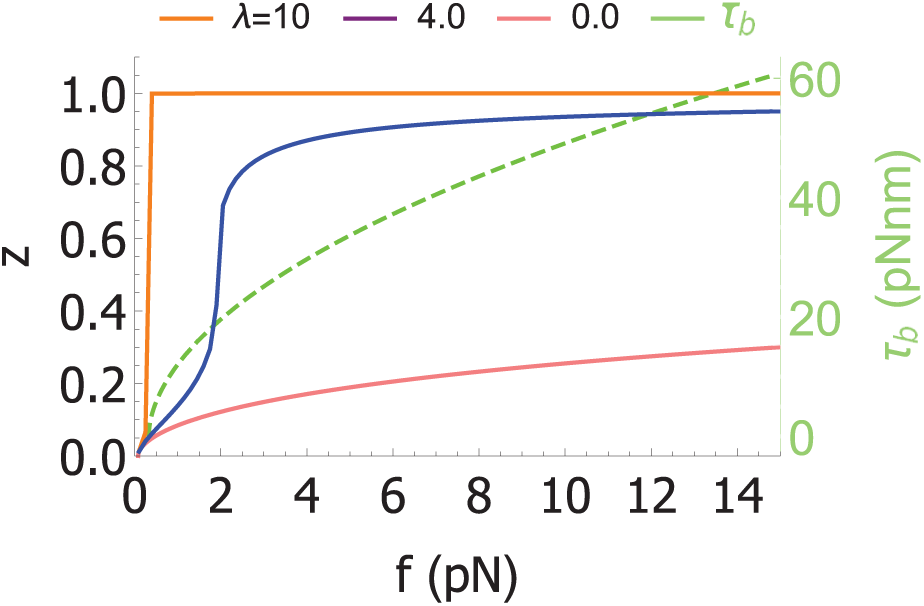
Collapse position *z*_*c*_ as a function of force for various fixed topoisomerase rates (top legend) showing how replication can be completed at lower forces and buckling torque *τ*_*b*_ (shown in green) with topoisomerase action.

**FIG. 4:**
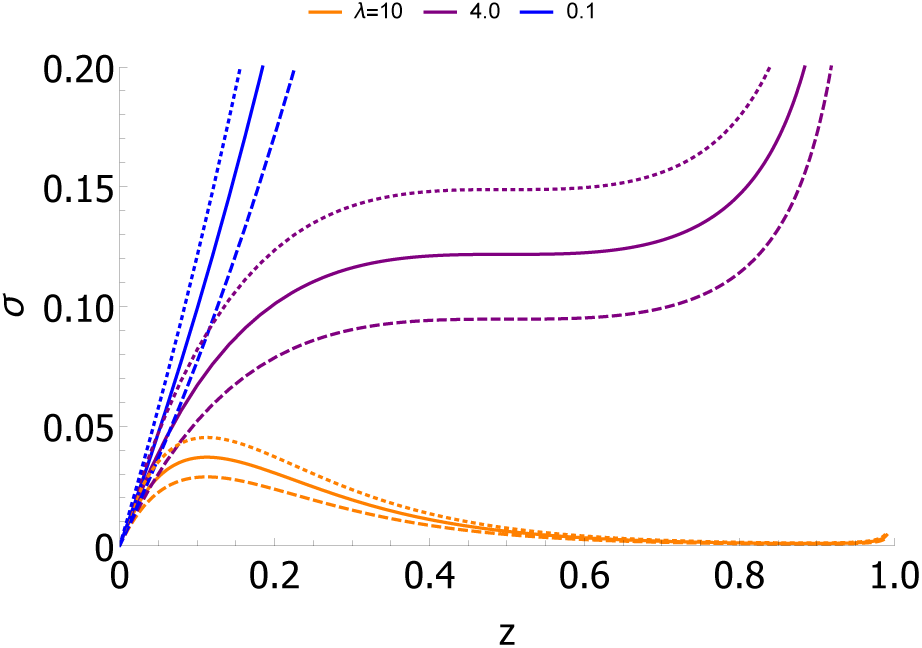
Super-coiling density *σ* as a function of fork position *z* = *x/L* against an fixed barrier (solid) and a long gene convergently (dashed) or divergently (dotted) oriented for various topoisomerase action rates (top legend) at fixed force *f* = 1*pN*. Depending on gene orientation SCD can be raised or lowered resulting in altered collapse position. The 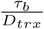 contribution is dropped and 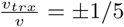 is used.

**FIG. 5:**
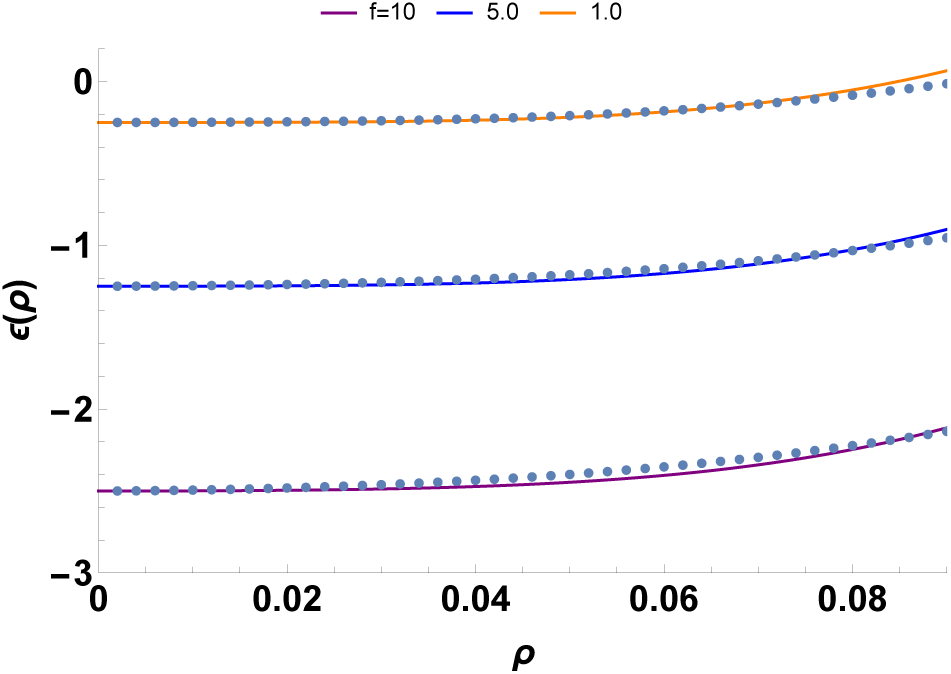
Comparison of the free energy given by phenomenological equation 29 to minimization of the full energy in equation 27 for a 0.1 *M* solution.

**FIG. 6:**
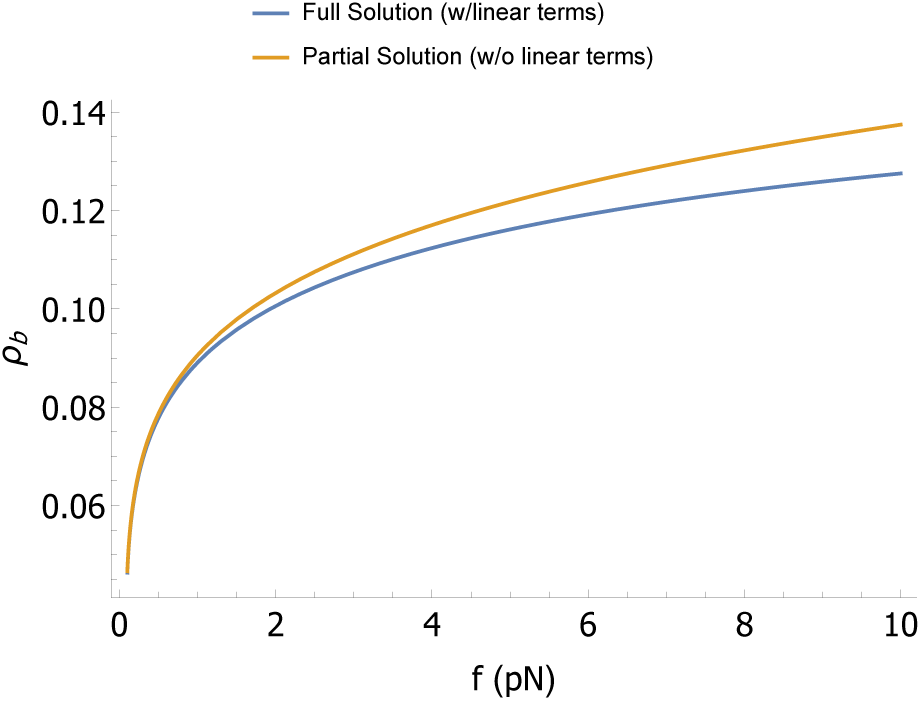
Buckled *ρ*_*b*_ comparison between the full solution given in the S.M. and partial solution used in main text.

During DNA replication additional topological barriers may exist to the free rotation of the unreplicated region presenting additional constraints for fork progression. A common obstruction to free DNA rotation is transcription and the interaction between fork progression and transcription can result in non-local interactions and stalling between transcription and replication [9]. We can incorporate the effect of an area of active transcription, and thus the absence of a static barrier, by changing the boundary condition at this point by introducing an addition degree of freedom for DNA *φ*. Then the super-coiling density in the unreplicated region (between the replication fork and the area of active transcription) is given by

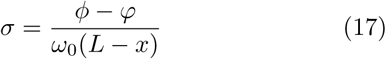

which leads to a modified equation 12

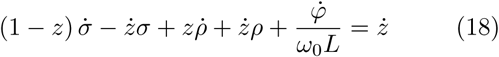

During transcription RNA polymerase can act as both a barrier to free DNA rotation as well as a source of SCD [17] leading to numerous interesting behaviors [18]. To incorporate this effect the dynamic mechanical response of *φ* is given as

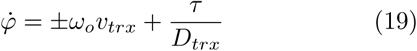

where *v*_*trx*_ incorporates the ability of the gene to inject more SCD in the region which is positive (negative) when the gene is convergently (divergently) oriented with the direction of replication and *D*_*trx*_ reflects the drag associated with difficulty in rotating the RNAP and nascent RNA (which may depend on the length of the gene).

The solution for equation 18 including non-linear terms is difficult, however in the buckled case the non-linear terms vanisand the action of topoisomerases can be included as before as before to find a modified equation 14

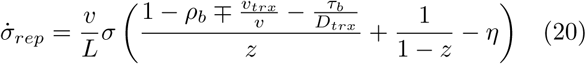

yielding a solution

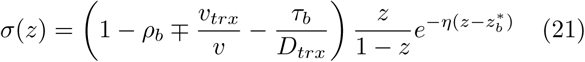

resulting in a modified equation 15. As before this solution has a local maximum provided the rates yield real solutions given by 16. However the slope in equation eq:sctrxdnysolution changes the effective buckling density, and thus the the ability for 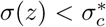 during replication, as a function of gene length and orientation allowing for varying amounts of replication to occur before collapse. This important difference is highlighted in figure 4 revealing the role of gene orientation in replication transcription conflicts. This effect could be responsible for additional phenomena related to transcriptional changes as a function of a gene’s relative position and orientation to replication.

Other barriers could be incorporated in this framework resulting in modified topological and mechanical constraints. However, the central framework of this work should remain valid. For example the presence of multiple replication forks can be accommodated resulting in modified equations for BD and SCDs as a function of total replication completed (See SM for details).

Many important aspects of DNA replication are not addressed here and the simple nature of the model developed in this letter leaves a number of opportunities for future work to bridge the divide between theory and experiment. To date, the basic elements of the topological and mechanical competition cells encounter during DNA replication and cellular division have remained mysterious. It is the aim of this work to take some of the first steps in constructing a full theoretical picture of this mechanical process and the basic features which cells must overcome.

In this vein we have neglected many important aspects of DNA replication that fall into two rough categories: behavior of the replication machinery during replication and the dynamic behavior of the braided and super-coiled DNA. In this first effort we have only been concerned with developing a model that accommodates the equilibrium behavior of braided, replicated and super-coiled, unreplicated DNA. Future models and experiments that incorporate the dynamical response of both regions may reveal important aspects of the process of DNA replication. Additionally, we have only evaluated the equilibrium response of DNA replication for linear DNA at constant forces. A substitution of a closed ring topological boundary condition (as would be expected for bacterial chromosome) instead of fixed force boundary conditions will change the torsional response of the unreplicated region. Finally including identifying the sources of the torque present during DNA replication may uncover the means by which helicase is able to unwind DNA as well as an observed dead-mans switch between the construction of new replicated strands and the progression of the replication fork [7].

The elements and properties of the idealized model of replication presented in this letter should be viewed as first steps in understanding the full mechanical nature of DNA replication. However, even in the simple framework presented here a number of important phenomena have been outlined, most notably the competition between topology and mechanics cells must overcome to succesfully replicate DNA. During this process fork progression and the resulting torsional load put onto DNA must be dealt with before cells can divide and the basic elements of this competition have been highlighted in this letter. Additionally, the basic properties of non-local interaction between replication and transcription have been outlined. This framework should serve as a foundation for future studies to bridge the divide between the simple properties of DNA replication and a more complete description needed to fully understand DNA replication in cells.

This work was supported by the National Science Foundation Center for Theoretical Biological Physics (Grant NSF PHY-1427654). S.A.S. thanks Herbert Levine, Edward Banigan, Hugo Brando and Sumitabha Brahmachari for helpful discussions.

## SM

### Phenomenological equations for braided DNA torsion

We would like to find simple equations for the equilibrium torsional response of braided DNA as a function of braid density (BD). This can be obtained by analyzing the free energy density for braided DNA at equilibrium. Using [15] the free energy per unit length of a polymer with braid radius *R* and pitch *P* held at a fixed force *f* is given by

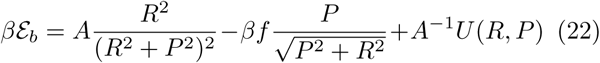

where *A* = 50*nm* is the bending persistence of double stranded DNA with *β* = 1*/k*_*b*_*T* and *k*_*b*_*T* = 4*pNnm* (at 290 K) throughout this analysis.

The electrostatic energy *U* (*R, P*) of the braid is given by

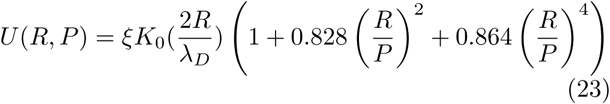

where *λ*_*D*_ is the Debye-screening length and *ξ* the amplitude of the Debye-Hückel potential, both of which depend on the chemical composition of the solution the braid is in [15]. We will use the electrostatic parameters corresponding to a salt concentration of 0.1 *M*

To write the energy eq.22 in terms of the braid density we can make the substitution *ρ* = *θ/*(*ω*_0_ *L*) where *θ* is the angle by which the two strands of length *L* have been braided and *ω*_0_ = 2*π/*(3.6*nm*) encodes the natural linking density of DNA. For a braid of fixed length *L* we can use need the geometric relationship 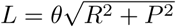 for the helical wrapping of the braid. This leads to the expression

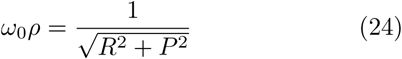

where *η* = 2*πω*_*o*_. Solving for the pitch yields

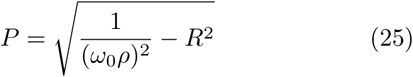

so that the energy per unit length is given by

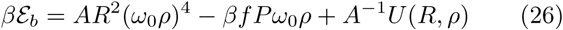

substituting the density constraint from eq.24 and expanding for small *ρ* gives

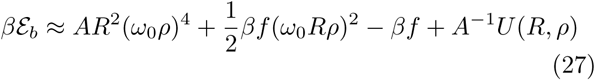

We can now minimize equation 26 with respect to *R* at fixed force and BD *ρ* yielding an effective energy density

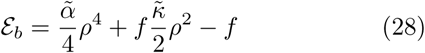

where 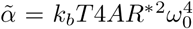 and 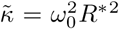. The resulting radius *R*^*^ after minimization has little *ρ* dependence and we have dropped the electrostatic contribution in the effective energy density B. This is done as when minimized it contributes little to the energy density serving mainly to set the braid radius *R*. This radius is set by the electrostatic contribution by the energy which is driven by short distance electrostatic interactions.

Comparisons between the full minimization of eq.22 to the phenomenological equation are shown in figure

Finally, the torsional response of braided DNA is given by the relationship

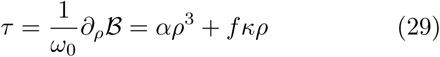

where for physiological conditions stated above the resulting values for the phenomenological equations are *α* = 13, 781 *pNnm* and *κ* = 7.7 *nm*. Since *α* ≫ *κ* and *ρ* ≪ 1 we can the linear term in the torsional analysis presented in the letter. A full solution to the SCD as a function of fork position including the linear *ρ* terms is given below.

### Comparison to full solution

It is possible obtain a solution for the BD during replication with linear *ρ* which were dropped in the analysis presented within the letter. To do this let us start from equation 7 of the main text but now using

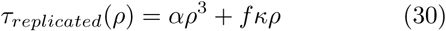

for the torsional response of the replicated region we find the below equation for the BD as a function of fork position before buckling

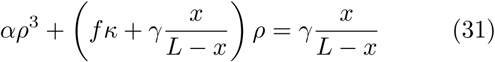

and after buckling

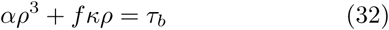

As discussed in the main text *τ*_*b*_ sets the torque in the system given by the buckled response of the unreplicated region. These equations yield a non-linear solution for the braid density as a function of replication position

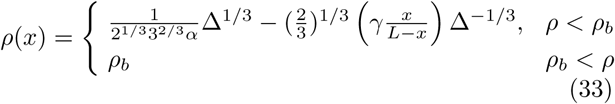

with

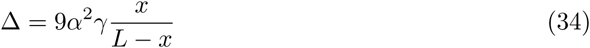

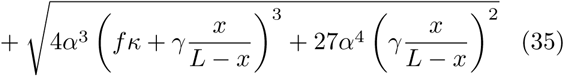

and

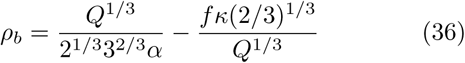

with

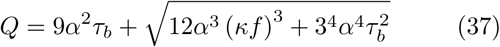

which is compared to the simplified expression for *ρ*_*b*_ used in the text in figure. Having obtained the BD, the solution for the SCD using the full solution follows in the same manner outlined in the main text.

### Multiple replication forks

Following the logical presented in the main text we can examine the behavior of multiple replication forks. For the case of *N* replication regions each of size *x*_*i*_ of total size 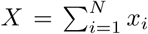 flanked by unreplicated regions of size *z*_*i*_ of total size 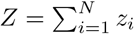 existing in a region of size *L* = *Z* + *X* (in any arrangement) a generalization to equation can be made to equation 2. First noticing that the same topological constraint must be true

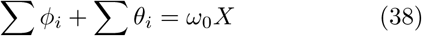

where *phi*_*i*_ is the twist in the unreplicated region on one side of the replicated regions which have braiding *θ*_*i*_. Yields a generalized equation 2

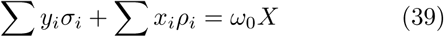

where the SC and braid densities in each region are given by *ω*_0_ *σ*_*i*_ = *ϕ*_*i*_*/z*_*i*_ and *ω*_0_ *ρ*_*i*_ = *θ*_*i*_*/x*_*i*_ respectively.

Since the entire DNA is connected we will take the torsion in the entire region *L* to be homogeneous. Thus we must have for each boundary between replicated and unreplicated regions *τ*_*rep*_ = *τ*_*unrep*_ and thus we must have *σ*_*i*_ = *σ*_*j*_ and *ρ*_*i*_ = *ρ*_*j*_ for all regions of unreplicated and replicated regions respectively. Equation 39 then becomes

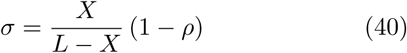

which is identical to the form taken by equation 2. Then agreement between the torsion in the replicated and unreplicated regions yields modified equations with *X* substituted for *x* thus identical equations for the braid and super-coiling density (eqs.9,11). Additional effects such as the role of topoisomerase could be added within this framework resulting in solutions closely following those presented in the main text for the case of one replication fork.

